# Silencing *XIST* on the future active X: Searching human and bovine preimplantation embryos for the repressor

**DOI:** 10.1101/2021.10.05.463256

**Authors:** Melis A Aksit, Bo Yu, Bernard AJ Roelen, Barbara R. Migeon

## Abstract

X inactivation is the means of equalizing the dosage of X chromosomal genes in male and female eutherian mammals, so that only one X is active in each cell. The *XIST* locus (*in cis*) on each additional X chromosome initiates the transcriptional silencing of that chromosome, making it an *inactive* X. How the *active* X in both males and females is protected from inactivation by its own *XIST* locus is not well understood in any mammal. Previous studies of autosomal duplications suggest that gene(s) on the short arm of human chromosome 19 repress *XIST* function on the active X. Here, we examine the time of transcription of some candidate genes in preimplantation embryos using single-cell RNA sequencing data from human embryos and qRT-PCR from bovine embryos. The candidate genes assayed are those transcribed from 19p13.3-13.2, which are widely expressed and can remodel chromatin. Our results confirm that *XIST* is expressed at low levels from the future active X in embryos of both sexes; they also show that the *XIST* locus is repressed in both sexes when pluripotency factors are being upregulated, during the 4-8 cell and morula stages in human and bovine embryos – well before the early blastocyst (E5) when *XIST* on the inactive X in females begins to be upregulated. Our data suggest a role for *DNMT1, UHRF1, SAFB* and *SAFB2* in *XIST* repression; they also exclude *XACT* and other 19p candidate genes and provide the transcriptional timing for some genes not previously assayed in human or bovine preimplantation embryos.

## INTRODUCTION

Only one X chromosome is active in diploid human cells, irrespective of the sex of the individual and the number of X chromosomes^1^. *XIST*, a non-coding RNA^2^ emanating from the X inactivation center, has been shown to be a potent chromosome silencer – not only for the inactive X in all the eutherian mammals studied^3-6^ but also for autosomal chromosomes into which it has been transfected^7,8^. Although another non-coding RNA, antisense to *XIST*, seems to have a role in protecting the mouse *active* X, its human counterpart, *TSIX*, was truncated during mammalian evolution^9,10^, is co-expressed with *XIST* from the inactive X in male and female cells ^10^ and has not been shown to play a role in protecting the active X in species other than rodents. Further, *TSIX* has not been found in many mammalian genomes. (https://genome.ucsc.edu/cgi-bin/hgGateway.6/26/21)

Human triploid cells (69, XXX and 69, XXY) have two *active* X chromosomes^11-17^ because a second X is protected from silencing by the extra set of autosomes in triploid cells. The simplest explanation is that active X’s are chosen by repressing their *XIST l*oci; the key repressor is encoded by an autosome ^11,18-20^.

Previously, we identified two candidate autosomes^11^ based on a comprehensive study of human trisomies (47,XX). We could exclude the X chromosome and all autosomes, (except 1 and 19), because unlike triploids, these 21 different trisomies had only one active X^11,14^. Trisomies 1 and 19 could not be studied, as they do not survive implantation, presumably because they are gene-dense. Therefore, we ascertained all available partial trisomies of chromosome 1 and 19 that *survive* gestation. By searching the literature to determine the trisomic regions that are tolerated in liveborns, we could exclude them from our candidate regions of chromosomes 1 and 19^11^. In 2017, we could analyze duplications of chromosome 1 and 19 in the *Decipher* database, which records the sex and phenotypes of chromosomal duplications, deletions and single nucleotide variants^19^. Because extra doses of repressor would not be lethal for males with only one X chromosome, yet cause the death of females with two active X chromosomes, we looked for *sex* differences in duplications on chromosomes 1 and 19; as controls, we performed the same search for deletions on chromosomes 1 and 19, and for duplications on all the other human autosomes in the *Decipher* database. What we observed was that only chromosome 19 had an extensive region (~ 8 MB) on its short arm (19p13.3–13.2), that is *intolerant* of interstitial or tandem duplication in females^21^, suggesting that it contains the key dose-sensitive gene(s) that induces *XIST* repression^19,21^. The sex difference in duplications on chromosome 19p was highly significant (<10^−11^) (based on a permutation test)^19^.

This region of chromosome 19 includes >290 genes, which code for proteins as well as epigenetic factors^19^. Many of these genes could be eliminated from consideration as they are expressed in single tissues, or are expressed only after implantation, or have known functions incompatible with *XIST* repression. The most relevant genes were the chromatin remodelers that were expressed in every tissue. We knew that epigenetic factors could repress *XIST in trans*. Of particular interest were the writers and erasers of epigenetic marks, such as the lysine demethylases, which have been implicated in *Xist activation* in mice^22,23^. If lysine demethylases could activate *Xist* expression, other epigenetic marks might prevent it, perhaps by histone changes leading to DNA methylation.

Among the relevant 19p candidate genes are the DNA methyltransferase, *DNMT1*, its co-factor *UHRF1*, the satellite attachment factors *SAFB1* and *SAFB2*, the hetero-ribonuclear protein *HNRNPM*, the histone methylases, *KDM4B*, and *KDT2B*, the long non-coding RNA (lncRNA) *TINCR*, as well as a cluster of genes coding for zinc finger proteins (Table 1). We analyzed these genes as they seemed able to repress *XIS*T.

**Table 1:** The genes analyzed in human preimplantation embryos.

| Gene |  | Function | Position<br>(5' end) | Expressed<sup>a</sup> |
| --- | --- | --- | --- | --- |
| *XIST* | Xq13.2 | Long non coding RNA | 73,820,650 | + |
| *SIRT6* | 19p13.3 | Histone deacetylase | 4,174,108 | + |
| *UHRF1* | 19p13.3 | Tethers DNMT1 to chromatin | 4,903,079 | + |
| *KDM4B* | 19p13.3 | XIST activator (ref) | 4,969,122 | - |
| *PTPRS* | 19p13.3 | Protein tyrosine phosphatase receptor | 5,205,502 | - |
| *ZNRF4* | 19p13.3 | Zinc finger, ring finger protein | 5,455,526 | - |
| *TINCR* | 19p13.3 | Long non-coding RNA | 5,558,166 | + |
| *SAFB2* | 19p13.3 | Scaffold attachment factor | 5,586,992 | + |
| *SAFB* | 19p13.3 | Scaffold attachment factor | 5,623,034 | + |
| *RFX2* | 19p13.3 | Methylation binding protein | 5,993,163 | - |
| *HNRNPM* | 19p13.2 | Heterogenous nuclear receptor protein M | 8,444,574 | + |
| *ZNF414* | 19p13.2 | Zinc finger protein 414 | 8,575,462 | - |
| *UBL5* | 19p13.2 | Ubiquitin like protein | 9,827,879 | + |
| *DNMT1* | 19p13.2 | DNA methylase | 10,133,343 | + |
| *SMARCA4* | 19p13.2 | SWI/SNF chromatin regulator | 10,960,932 | + |
| *PRMT4* | 19p13.2 | Arginine methyltransferase | 10,982,189 | - |
| *ZNF823* | 19p13.2 | Zinc finger protein | 11,832,080 | + |
| *ZNF69* | 19p13.2 | Zinc finger protein | 11,887,772 | - |
| *XACT* | Xq23 | Long non coding RNA | 113,616,300 | - |
| *NANOG* | 12p13.21 | Homeobox transcription factor | 7,781,401 | + |
| *POU5F1* | 6p21.33 | Pou domain transcription factor | 31,164,336 | + |
| *SOX2* | 3q26.33 | Neural inhibitory factor | 181,711,924 | + |

<sup>a</sup>. Genes with an average expression above 5 RPKM at any stage in either dataset are considered expressed.

Previous studies showed that *XIST* is expressed – albeit at levels that cannot silence the chromosome – from every X in human males and females at the four to eight cell stage ^24^. Silencing of X chromosomal genes occurs only when *XIST* is upregulated from the future *inactive* X, beginning at the early blastocyst stage^25,26^. Because the *XIST* locus on the future *active* X needs to be silenced before *XIST* is upregulated on the future *inactive* X, we ascertained the time during preimplantation development when our candidate genes are transcribed. We examined RNA transcripts collected from preimplantation human embryos, using the single-cell RNA-sequencing data present in the two published datasets^25,26^ available to us. Analyzing their expression, we identified candidate genes with patterns that suggest they could be involved in *XIST* repression.

We validated the human expression patterns in bovine preimplantation embryos. It has been shown that the bovine is a *more appropriate* model for human X-inactivation than the mouse^3^. Both human and cattle are mono-ovulatory and the timing of oocyte development and embryonic genome activation as well as the embryonic transcriptomes are more similar than they are to mouse^27^. Bovine *XIST* is also not repressed by its antisense gene, as *TSIX* has not been found in the bovine genome *(https://genome.ucsc.edu/cgi-bin/hgTracks?db=bosTau9*). Bovine, like humans and many other mammals except rodents, have conserved this block of candidate genes over millions of years of evolution and the chromosome 19p genes are clustered on bovine chromosome 7 ^28^.

Because of the many species differences in development^18^, we did not expect the bovine to regulate X chromosomes exactly like humans, but *unlike rodents* they do not imprint X inactivation, they conserve the 19p gene cluster on bovine chromosome 7 and they do not use *TSIX* to protect the active X.

## MATERIALS AND METHODS

### Single-cell RNA Sequencing Datasets

Data from single cell RNA-sequencing (scRNA-seq) of human preimplantation embryos were taken from two publicly available datasets: *Dataset 1*: Yan, et al. (GEO accession: GSE36552)^26^ and *Dataset 2*: Petropoulos, et al (EMBL-EBI accession: E-MTAB-3929)^25^. See **Supplemental Table 1** for specimens that are included in each dataset. These two datasets available for our analysis have different content of embryos.

### Sexing of Human Embryos using scRNA-seq

Sexing of embryos of *Dataset 1*^26^ was conducted by Moreira de Mello, et al (2017)^29^ using Y-linked expressed genes outside of the pseudoautosomal region; sexing begins at the 8-cell stage (Supplemental Table 1). Sexing in *Dataset 2* was conducted by Petropoulos, et al (2016)^25^ (Reinius, B, personal communication), also based on the expression of the Y-linked genes outside of the pseudoautosomal region as described in Petropoulos, et al (2016)^25^.

### Bovine in vitro embryo production and sample collections

Bovine ovaries, collected from a local slaughterhouse, were transported to the laboratory in a thermos flask. After a water rinse at 30°C, the ovaries were kept in 0.9% NaCl supplemented with penicillin/streptomycin (100 µg/mL) at 30 °C. Cumulus-oocyte complexes (COCs), aspirated from follicles with a diameter of 2-8mm, were identified using a stereomicroscope. The COCs were matured *in vitro* and fertilized as described previously^30^. After 23h maturation, COCs were transferred to fertilization medium and incubated with 10^6^/mL sperm cells. Female or male embryos were derived from X or Y-sorted sperm (CRV, Arnhem, the Netherlands)^3^. The moment that sperm were introduced into fertilization medium was considered day 0. After incubation with sperm for 20-22h, presumptive zygotes were freed from cumulus cells by vortexing for 3 min, and placed in synthetic oviductal fluid (SOF) in a humidified atmosphere at 5% CO_2_ and 7% O_2_ at 39 °C1. At day 5, cleaving embryos were transferred to fresh SOF and further cultured until day 8.

For subsequent analysis, zygotes, two, four and eight cell embryos were collected at 20, 32, 38 and 56 h after the start of fertilization, respectively. Morulae and blastocysts were collected on day 5 and day 8, respectively. Embryos in pools of 20 were stored in RLT buffer (Qiagen, Valencia, CA, USA) at −80 °C until RNA extraction (*see Supplementary Experimental Data*).

## RESULTS

Expression levels of *XIST*, candidate *XIST* repressor genes and three zygotic activation marker genes (*NANOG, POU5F1*, and *SOX2*) were evaluated in two published human single-cell RNA-seq datasets (**Table 1**), to determine when *XIST* repression occurred, and which candidate genes were expressed at that time. Our candidate genes included genes from the region of chromosome 19p we previously demonstrated has a sex bias^19^. *NANOG, POU5F1*, and *SOX2* were studied because they are known to be upregulated immediately after transcription transitions from maternal oocyte to zygote in human pre-implantation embryos^31^. Also we studied the primate specific X-linked lncRNA, *XACT*, as others suggest it is the gene that protects the active human X from being silenced by its *XIST* locus^32^. Because 47, XXX diploid cells with three copies of *XACT*, have only a single active X, we do not consider *XACT or any other X-linked gene to be* a serious candidate for the *XIST* repressor (see **Discussion**); however, we wished to determine if *XACT* was expressed at the time of *XIST* repression. Whenever possible, we used the sex of the embryo to determine if expression of the locus occurred in males as well as females.

*XIST* was first expressed at extremely low levels in the 8-cell or morula stages by both male and female embryos, but it showed a female sex bias in the early blastocyst stage (**Figure 1**). In *Dataset 1*^26^ *XIST* was not expressed until the 8-cell stage (**Figure 1A**; Oocyte, Zygote, 2-cell and 4-cell average expression values: 0 RPKM). *XIST* expression was very low in both sexes at the 8-cell stage; it increased in the morula and was greatest at the late blastocyst stages of female embryos (**Figure 1A**; 8-cell expression: 0.48 RPKM, morula (E4) average expression: 1.01 RPKM, early blastocyst (E5) average expression: 3.31 RPKM). In *Dataset 2*^25^ the expression of *XIST* was also very low at the 8-cell stage (0.09 RPKM); it increased from morula (E4) to E5-E7 blastocyst stages (**Figure 1A**, average expression 2.91, 4.25, 11.38 and 9.09 RPKM, in E4, E5, E6 and E7, respectively). It is clear from *Dataset 2* that female embryos are the ones that express *XIST* in the early (E5) blastocyst (average female *XIST* expression: 6.61 RPKM; average male *XIST* expression: 0.67 RPKM), confirming that this is the time when *XIST* begins to be up-regulated in females – eventually silencing the future inactive X chromosomes.

**Figure 1:**
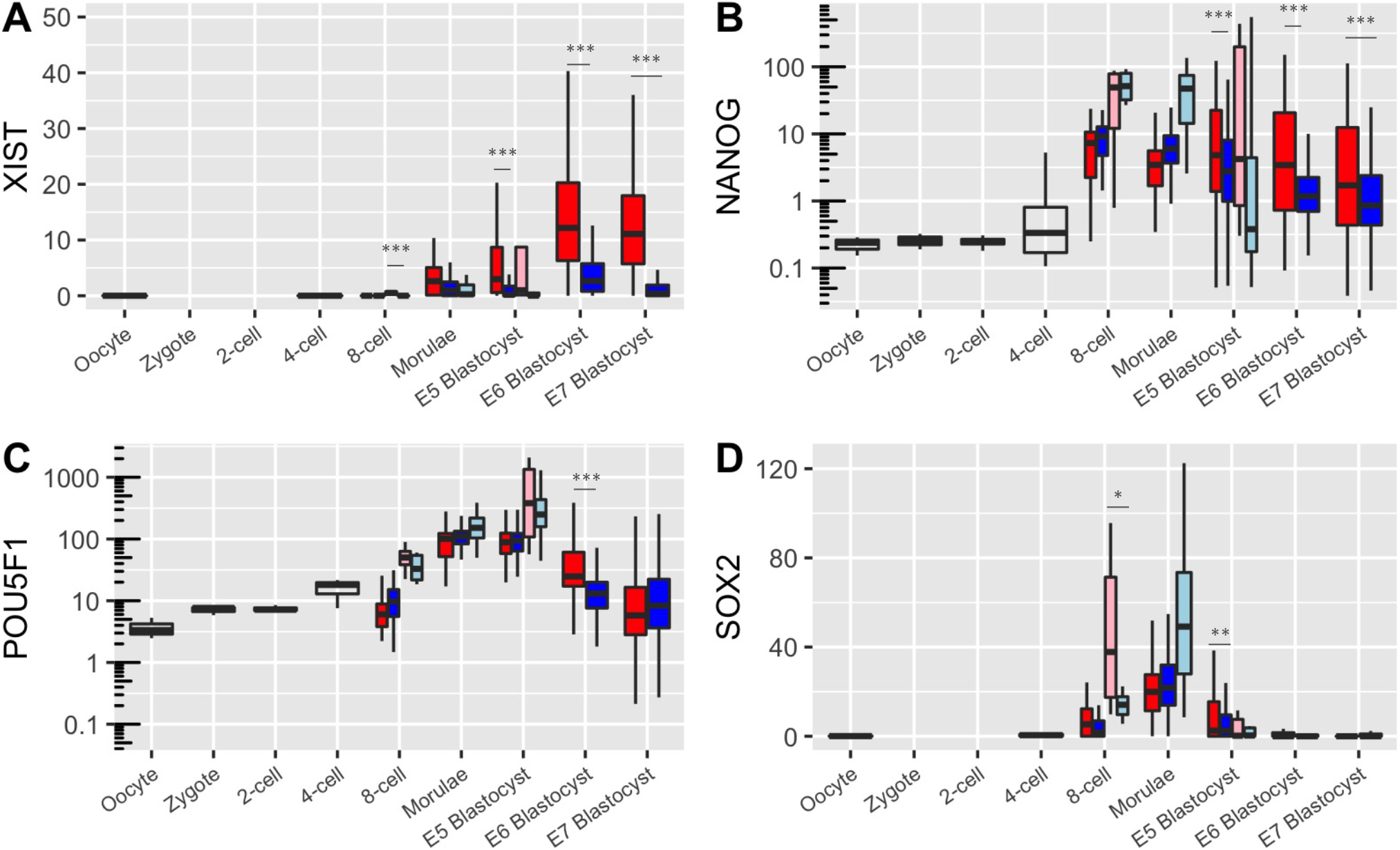
Box plots of gene expression (RPKM) *of XIST, NANOG, POU5F1* and *SOX2* determined by scRNA-seq in preimplantation human embryos. *Embryos are grouped by sex and dataset (Pink: Females from dataset 1, Light blue: Males from dataset 1, No fill: undetermined sex from dataset 1, Red: Females from dataset 2, Blue: Males from dataset 2). The horizontal line in the middle indicates median value, and the boxes show the first and third quartiles. The upper whisker extends from the hinge to the largest value no further than 1*.*5 * IQR from the hinge (IQR: inter-quartile range; distance between the first and third quartiles). The lower whisker extends from the hinge to the smallest value at most 1*.*5 * IQR of the hinge. Data beyond the end of the whiskers are not plotted. Significant differences between females and males are indicated by * (p < 0*.*05), ** (p < 0*.*01) and *** (p < 0*.*005)*.

Figure 1 also shows the expression of *NANOG, POUF51* and *SOX2*, that are critical for maintaining pluripotency and are expressed at the time when tissues begin to differentiate^33,34^.. The first two genes were not detectable until the human eight-cell – morula stages, reaching a maximum in the early blastocyst (E5). *SOX2* was transcribed at the eight-cell stage and is most abundant in the morula. Females expressed *NANOG* at higher levels at the E5-E7 blastocyst stages. Although the expression values vary between the two datasets due to technical variance associated with the sequencing technology and between sexes due to individual variation, whether or not the gene is expressed at the various stages is consistent.

Because 47, XXX females with three copies of the X-linked lncRNA, *XACT*, have two inactive X’s and only one active X, we had previously eliminated *XACT* and other X-linked genes as candidate *XIST* repressors. And in our analysis, we did not find that *XACT* was expressed in Dataset 1 (**Table 1**). Petropolis et al^25^ (Dataset 2) reported its expression at very low levels (<3 RPKM) in morulae (E4) and E5-E7 blastocysts, which is below our cut-off for being expressed (5 RPKM) (**Table 1** (see Figure S5 in Petropoulos et al^25^). *XACT* may function mainly in maternal oocytes where it has been observed to be expressed from both X chromosomes in human primordial germ cells, but not in ovarian somatic cells^35^; it has been proposed to play a role in imprinting maternal genes in primordial germ cells^35^. Our interpretation that *XACT* has no role in protecting the active X is supported by recent studies of human embryonic stem cells that indicate that the deletion of *XACT* has no effect on X inactivation, but does perturb neuronal development in those cells^36^. The evidence, including this paper, is now definitive enough to eliminate *XACT* as a candidate *XIST* repressor.

### Candidate genes that could be eliminated from consideration because they are not expressed during pre-implantation

Among genes that were not transcribed in the preimplantation human embryo are *KDM4B, PTPRS, ZNRF4, ZNF414, RFX2* and *PRMT4*, and so these genes can be eliminated from our list of candidates (**Table 1**;**Supplemental Figure 1**). *SIRT6* and *TINCR*, which are not expressed prior to *XIST* upregulation, are also unlikely to be key *XIST* repressors (**Supplemental Figure 1**).

### Single-cell RNA-sequencing of human embryos shows that *DNMT1* and *UHRF1* are the candidate genes most highly transcribed prior to *XIST* upregulation

Of the genes evaluated (**Table 1**), the DNA methylase, *DNMT1* and its co-factor, *UHRF1*, were expressed the earliest and at the highest levels – paradoxically, at a time when DNA *demethylation* of the zygotic genome is prominent^37^. In *Dataset 1*, at the oocyte-4 cell stages, *DNMT1* was highly transcribed (averaging 1796.8, 2460.2, 4260.0, and 2440.1 RPKM in oocyte, zygote, 2-cell and 4-cell stages, respectively) when *XIST* was not expressed. In contrast, the levels of *DNMT1* transcription drastically decreased when *XIST* is upregulated (averaging 469.6, 301.3, and 4.2 RPKM, in the 8-cell, morula and late blastocyst stages, respectively), which was significantly different from the 4-cell stage (t-test p-values: 3.4e-13, 1.6e-13 and 1.5e-10 for 8-cell, morula and late blastocyst stages, respectively). *Dataset 2* showed the same trends; *DNMT1* was highly expressed at the 8-cell stage (mean 1005.6 RPKM), whereas its expression was much less at the morula-blastocyst stages (**Figure 2C**).

**Figure 2:**
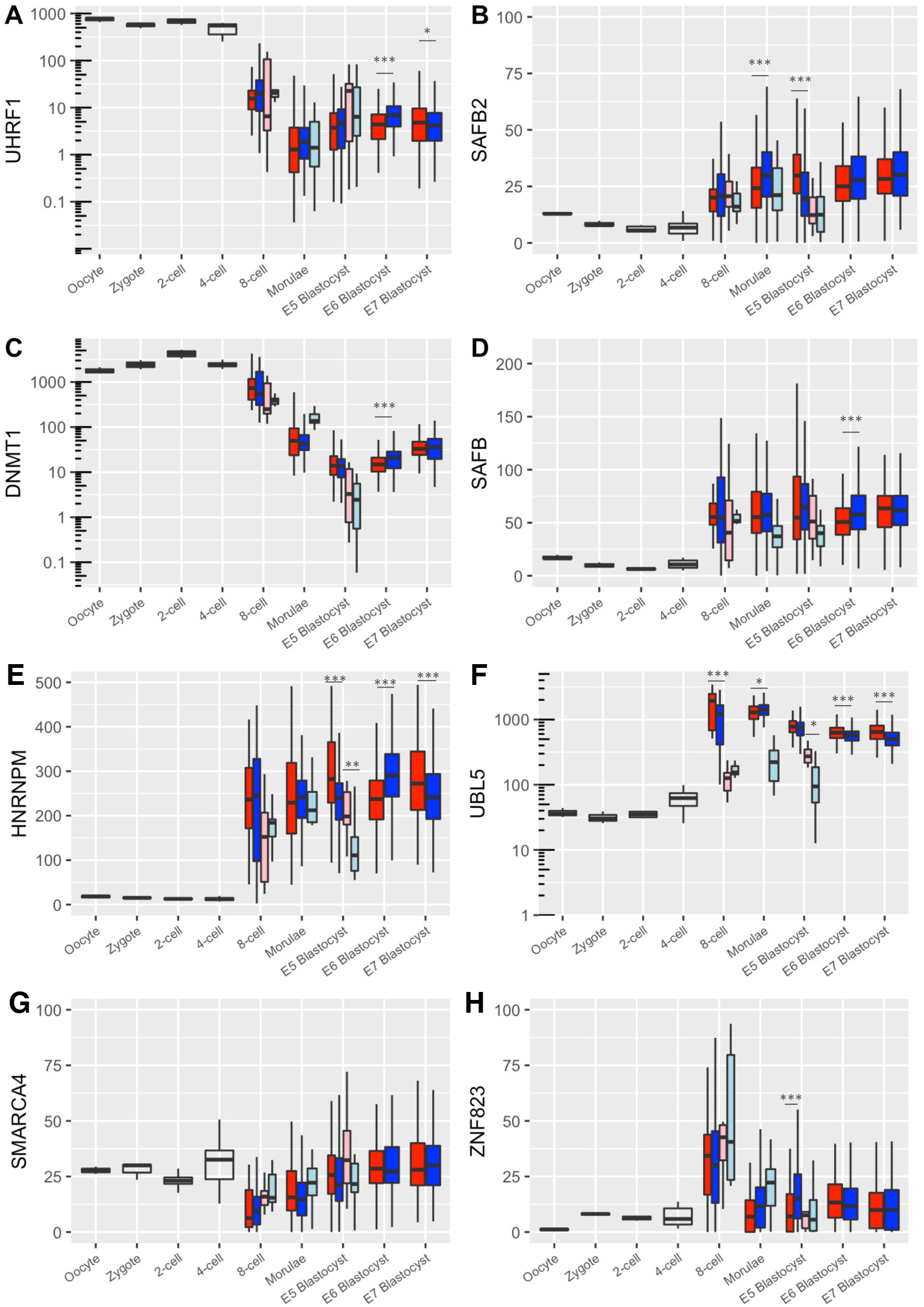
Box plots of gene expression (RPKM) for *UHRF1, SAFB2, DNMT1, SAFB, HNRNPM, UBL5, SMARCA4* and *ZNF823*, determined by scRNA-seq in preimplantation human embryos. *Significant differences between females and males are indicated by * (p < 0*.*05), ** (p < 0*.*01) and *** (p < 0*.*005)*.

*UHRF1*, coding for the DNMT1 co-factor had a similar expression pattern. For *Dataset 1* the average expression levels were 759.9, 560.1, 685.6 and 487.1 RPKM in the oocyte, zygote, 2-cell and 4-cell stages, respectively, significantly decreasing to 35.4, 3.1, and 18.7 RPKM in the 8-cell, morulae and late blastocyst stages, respectively (t-test p-values in comparison to the 4-cell stage: 6.3e-8, 8.7e-8, 1.1e-7, respectively). The results were similar for *Dataset 2* (**Figure 2A**).

### Other candidate genes transcribed prior to *XIST* upregulation

Albeit at a lower level than *DNMT1* and *UHRF1*, the genes for the satellite attachment factors *SAFB* and *SAFB2* were expressed prior to *XIST* upregulation (**Figures 2**); therefore, they remain candidates for the key *XIST* repressor. In addition, other genes in the region that encode nuclear proteins, *HNRNPM* and *URL5* were also expressed prior to *XIST* up-regulation (**Figure 2**). On the other hand, *SMARCA4*, which was transcribed prior to *XIST* upregulation (**Figure 2**) encodes a catalytic subunit of SWI/SNF complexes that remodels chromatin, making it more accessible to transcriptional activation – hence despite its transcription pattern, *SMARCA4* is less likely to be the XIST repressor. Though several zinc finger proteins were not transcribed in the early embryo, *ZNF823* was expressed during the four – eight cell stage (**Figure 2**).

### Bovine embryos show patterns of gene expression similar to humans

Because the patterns of *XIST* expression and silencing of the inactive X in bovine embryos are similar to those of human embryos^3,38^ we analyzed the expression of our candidate *XIST* repressor genes in bovine preimplantation embryos. The availability of sexed semen enabled us to generate sex-specific embryos. To confirm the accuracy of the sex-sorted sperm, we examined the expression of *DDX3Y*, which is transcribed from the Y chromosome in male embryos. As expected, the expression of *DDX3Y* was significantly higher in male than female embryos. The low level of expression in female embryos reflects the ~ 90% accuracy of the sperm sexing (**Figure 3**).

**Figure 3:**
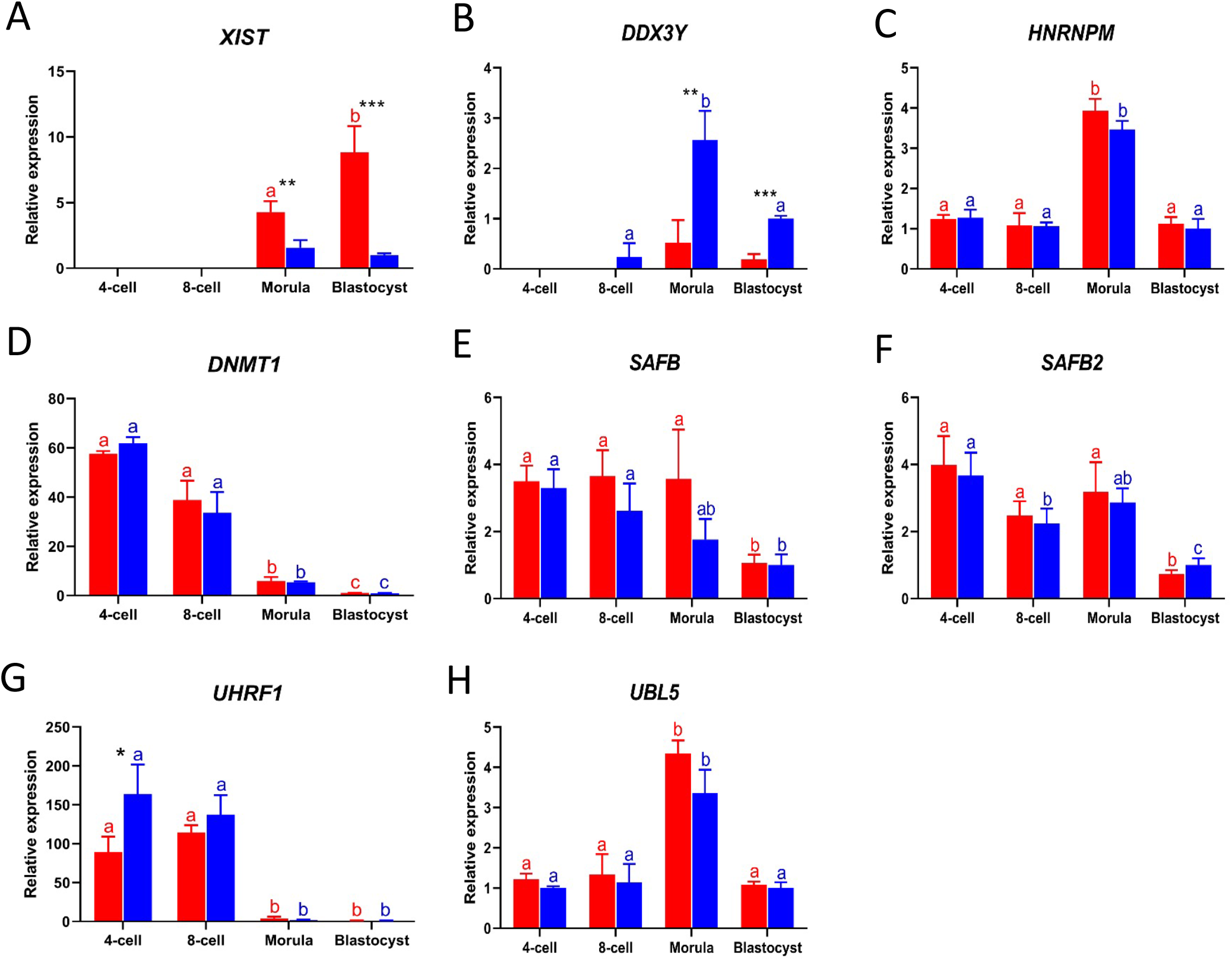
The relative expressions of candidate *XIST* repressor genes in female (red bars) and male (blue bars) bovine embryos from 4-cell stage to day 8 blastocysts, as detected by quantitative RT-PCR. *Embryos were derived by fertilization with sex-sorted sperm. Relative expression from male blastocysts set at 1. Significant differences between females and males are indicated by * (p < 0*.*05), ** (p < 0*.*01) and *** (p < 0*.*005). Significant differences among embryos with same gender are indicated by different letters with the same color (p < 0*.*05). Error bars indicate standard deviations of three independent biological replicates*.

Focusing on the expression of *XIST, DNMT1* and *UHRF1*, we found patterns similar to our human data. In unsexed bovine embryos, *XIST* expression has been detected at very low levels from the oocyte stage onwards, representing maternal transcripts^3^. The zygotic expression of *XIST* begins at the morula stage in both sexes and increases significantly at blastocyst stages only in female embryos (**Figure 3A**).

We detected the expression of *DNMT1* and *UHRF1* from the zygote stage onwards, at similar levels in male and female embryos (**Figure 3D**). Like in human embryos, *UHRF1* and *DNMT1* expression in the bovine embryos is downregulated after activation of the zygotic genome. Because zygotic activation occurs slightly later in bovine, downregulation of *UHRF1* and *DNMT1* also occurs slightly later in bovine **(Figure 3**). The expression of bovine *UHRF1* was enigmatically greater in male than female four-cell embryos (**Figure 3G**).

We also examined the expression of other candidate *XIST* repressors, *HNRNPM, SAFB, SAFB2* and *UBL5*, clustered on the region of bovine chromosome 7 that is homologous to the short arm of human chromosome 19. These genes were expressed throughout early preimplantation development, particularly in the morula (**Figure 3**).

## DISCUSSION

*XIST*, the long non-coding RNA is critical for silencing the *inactive* X. However, *XIST* needs to be silenced on the *active* X, so that the chromosome can continue to be transcribed and the promoter of the silent *XIST* locus on the active X is normally methylated in postnatal cells^39^. We have previously identified a region on chromosome 19 that we propose is involved in maintaining the transcriptional activity of the active X. Our comprehensive analysis of trisomies^11,19^ has directed us to that region on chromosome 19 and eliminated all other chromosomes, including the X, as the source of the *key XIST* repressor. Therefore, we hypothesized that some of the genes within the relevant region of chromosome 19, specifically those that could modulate chromatin, might be candidates for the key *XIST* repressor. Now, we have carried out an in-depth analysis of expression levels of some of the most likely candidate genes in this region during preimplantation development in both human and bovine embryos.

Our studies reveal that some of our human candidate genes are transcribed at the 4-8 cell stage (**Figures 2 and 3**) – the time that zygotic transcription occurs in the human embryo^31^. Bovine candidate genes are also expressed at the time of zygotic transcription, which occurs from the 8 cell-morula stages. This major transcriptional reprogramming event requires prior remodeling of the zygotic chromatin for transcriptional competence, and elimination of maternal transcripts. We propose that this is also when repression of the *XIST* locus on the future active X must occur; our observations suggest that it occurs with the upregulation of the pluripotency factors – the time when fetal developmental programs begin.

All the genes that remain candidates are involved in the regulation of chromatin. *HNRNPM, SAFB, SAFB2*, and *URL5* code for proteins which regulate transcription. Products from both *HNRNPM* and *UBL5* do not associate with protein; they only bind to RNA^40^ and have been implicated in the splicing of RNA. *XIST* has different splice variants^2^, but their importance to its function remains to be understood. Heterogeneous nuclear ribonucleoproteins, which directly bind to nascent RNA polymerase II transcripts, play an important role in processing heterogeneous nuclear RNAs to form mature mRNAs and in regulating gene expression. Ubiquitination is at the center of these mechanisms – occurring on all types of histones and regulating most nuclear signaling pathways.

The products of *UHRF1* and *DNMT1* place methyl groups on CPGs in the promoter region of genes, thus repressing their transcription. *UHRF1* codes for an E3 ligase, which is recognized by DNMT1, and their interaction stimulates the methyltransferase activity of DNMT1 – the key step in the maintenance of methylation patterns. Although DNMT1 is a maintenance methylase, it is also known to have *de-novo* methylating activity in the oocyte and early embryo^41,42^, and it is the imprinter of parentally imprinted genes^25^. In previous studies when *Dnmt1* is knocked out in male mice, *Xist* is expressed from the active X chromosome, indicating that *DNMT1* can repress *XIST*^39^.

The SAFB binding proteins are known to be satellite attachment factors. Both SAFB binding proteins, whose genes are in head-to-head orientation on human chromosome 19p13.2, have been shown to bind *XIST* as well as other non-coding RNAs^43^, but the consequences of that binding are not yet known.

The SAFB proteins, *UHRF1* and *DNMT1* and other candidates are transcribed in both sexes, as expected, because both males and females need to repress *XIST* on their active X. Because zygotic activation occurs after the 4 cell stage, the earlier transcripts from *UHRF1, DNMT1, HNRNPM, UBL5* and the *SAFB* loci in oocytes and zygotes, must be maternal in origin (**Figures 2 and 3**). Maternal RNAs are known to imprint some embryonic genes during oogenesis, including the maternal *Xist* gene in mice^35^. Yet, these genes must function in the fetal genome as well, as it is the duplication of 19p in the *fetal* genome that interferes with development of the female fetus. At the 4-8-cell stages, clearly these transcripts must originate from the fetus (**Figures 2 and 3**). Unfortunately, we have no means to determine from the present data if the origin of these gene transcripts at any stage is maternal or fetal or a combination of the two. Conceivably, transcription of *DNMT1* and *URHF1* begins in the oocyte and continues in the fetus after zygotic activation, and it is the fetal transcripts that are responsible for *XIST* repression.

Despite being >5MB apart on all the orthologs of chromosome 19p in a large number of mammalian species, the genes for methyltransferase DNMT1 and its co-factor UHRF1 remain together on that chromosome throughout millions of years of evolution^28^. Therefore, because they are transcribed from the pre-implantation embryo at the time of zygotic transcriptional activation, they are prime candidates for repression of *XIST* on the active X. This is also true for *XIST* binders *SAFB* and *SAFB2*. Of interest, the only other satellite attachment factor, *SAFA*, which is encoded by human chromosome 1, has been shown to be involved in silencing the *inactive* X by binding to mouse *Xist*^44^.

Our studies of the transcription of 19p genes DNMT1, URHF1 and several other 19pgenes that could repress the *XIST* locus on the active X of males and females reveal similar patterns of transcription in human and bovine, for considering the species differences in time of zygote activation. These studies provide further evidence that bovine is a good model for human X inactivation. As expected, the patterns are not identical, because of species variations that occurs throughout mammalian evolution. However, they suggest that transcription of the key *XIST* repressor may start in the oocyte, but that it is during the four to eight cell stage and morula when zygotic transcription begins, that *XIST* on the future active X is repressed in both males and females. Genes coding for *DNMT1*, its co-factor *UHRF1*, the satellite attachment factors, *SAFB* and *SAFB2*, the RNA binding proteins, *HNRNPM* and *URL5*, and zinc finger protein *ZNF823*, all residing on human chromosome 19p and bovine chromosome 7 remain candidates for the key *XIST* repressor.

Our results suggest that several of our candidate genes act together to repress the *XIST* locus on the future active X; they also provide a list of candidate genes that can be tested for their ability to repress *XIST* by knocking them out at the appropriate stage in human or bovine preimplantation embryos, or other appropriate non-rodent models. We also show that *XACT* is not the human *XIST* repressor, not only because of very low or non-expression in preimplantation embryos, but because it is X-linked.

For most eutherian mammals, the transcriptional silencing of *XIST* on the future active X provides an alternative model for X dosage compensation. X inactivation does not choose the inactive X, but chooses the X that will remain active by silencing its *XIST* locus. There is no need to count X chromosomes, as the future active X is chosen by silencing one *XIST* locus in both sexes, no matter the number of X chromosomes in the cell. In males, most often that X is their only X. Based on our studies, in human females, proximity to chromosome 19p plays a role in the choice. We know little about the nature of *trans*-interactions between chromosomes. Future studies should reveal how this is accomplished.

## Contents of Supplemental Material

2 Supplemental Tables and 2 Supplemental Figures

Supplemental Methods

## Acknowledgements

The authors are grateful to our Hopkins colleagues, Michael Beer, Hans Bjornsson, Haig Kazazian, Garry Cutting, Jeremy Nathans for their careful reading of the paper and their insightful suggestions. We also thank Drs. Bjorn Reinius and Frederik Lanner for their help with the sexing of embryos.

## Supplementary Methods

### RNA extraction, cDNA synthesis and qRT-PCR

Total RNA was extracted using an RNeasy Micro Kit (Qiagen, Valencia, CA, USA) and complementary DNA (cDNA) synthesis was performed ^3^. Primer sets (*Eurogentec*) were designed on Primer-Blast (http://www.ncbi.nlm.nih.gov/tools/primer-blast) using *Bos taurus* nucleotide sequences (Genbank; http://www.ncbi.nlm.nih.gov/nucleotide) as the template. To determine primer specificity and optimal annealing temperature, temperature gradients ranging from 55°C to 68°C were tested, using cDNA from 100 blastocysts. All amplification reactions were performed on three independent cDNA samples in duplicate, following the manufacturer’s protocol for the CFX detection system (*Biorad*). The reaction mixture was prepared with 10 µL iQ SYBR Green supermix (*Biorad*), 9 µL RNAse- and DNAse-free water (Invitrogen) and 1 µL cDNA with a final primer concentration of 500 nM. Reactions were started at 95 °C for 3 min, followed by 40 cycles of subsequently 95 °C for 20 sec, the primer specific annealing temperature (supplemental Table 2) for 20 sec and extension at 72 °C for 20 sec. To verify the purity of the PCR products, melting curves were plotted (with temperature increments of 0.5 °C) from 65 °C to 95 °C (each step for 5 sec). To determine quantitative (q) RT-PCR amplification efficiency, standard curves for each primer pair were made by 4 fold dilutions of cDNA from 400 oocytes or 100 blastocysts. Data were normalized using *GAPDH, SDHA* and *YWHAZ* as reference genes^45^.

### Statistical analysis

Statistical analysis of data from qPCR were carried out in Microsoft Excel and statistical differences were examined using GraphPad Prism 7 (https://www.graphpad.com/scientific-software/prism/). Differences between two groups were determined by two-tailed unpaired Students□t-tests and differences between multiple groups were analyzed by one-way ANOVA, followed by a post-hoc Tukey test. Statistical significance was set at P < 0.05. For human scRNA-seq statistical tests comparing gene expression over time, two-sample t-tests with unequal variance were used.

---

**Supplementary Table 1:** Distribution of embryos and cells in various stages in data obtained by Petropoulos, et al. and Yan, et al.

| Stage | Sex | Yan et al<br>Dataset 1 |  | Petropoulos et al<br>Dataset 2 |  |
| --- | --- | --- | --- | --- | --- |
|  |  | # of<br>embryos | # of<br>cells | # of<br>embryos | # of<br>cells |
| Oocyte | Unknown | 3 | 3 | - | - |
| Zygote | Unknown | 3 | 3 | - | - |
| 2-cell | Unknown | 3 | 6 | - | - |
| 4-cell | Unknown | 3 | 12 | - | - |
| E3 | Male | 1 | 8 | 7 | 48 |
| 8-cell | Female | 2 | 12 | 6 | 33 |
| E4 | Male | 2 | 16 | 7 | 98 |
| Morula | Female | - | - | 9 | 92 |
| E5 | Male | 2 | 22 | 14 | 206 |
| Blastocyst | Female | 1 | 8 | 9 | 161 |
|  | Female, X0 | - | - | 1 | 10 |
| E6 | Male | - | - | 8 | 180 |
| Blastocyst | Female | - | - | 10 | 235 |
| E7 | Male | - | - | 10 | 176 |
| Blastocyst | Female | - | - | 7 | 290 |

**Supplementary Table 2:**
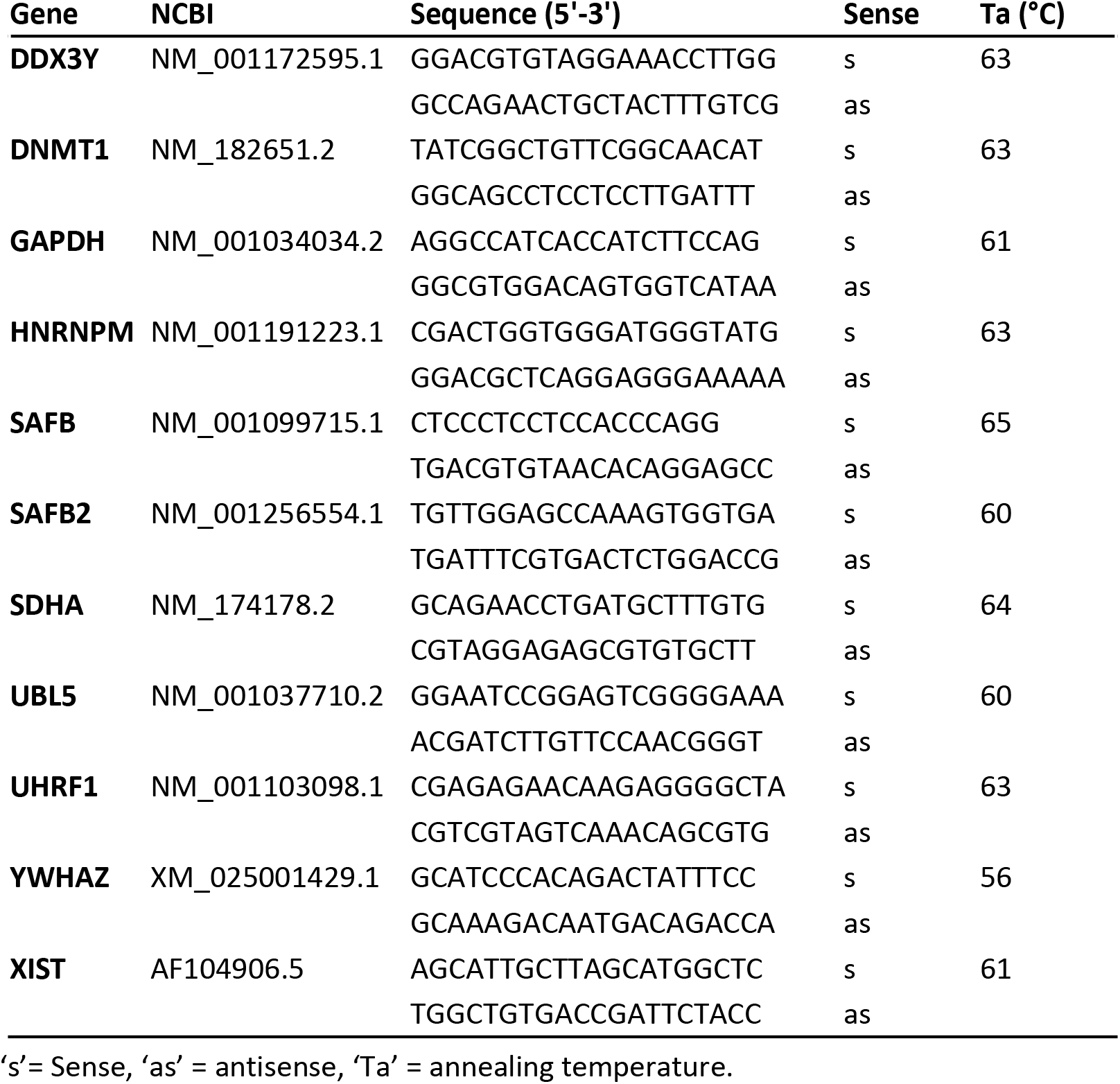
Primers used for quantitative RT-PCR analysis

**Supplemental Figure 1:**
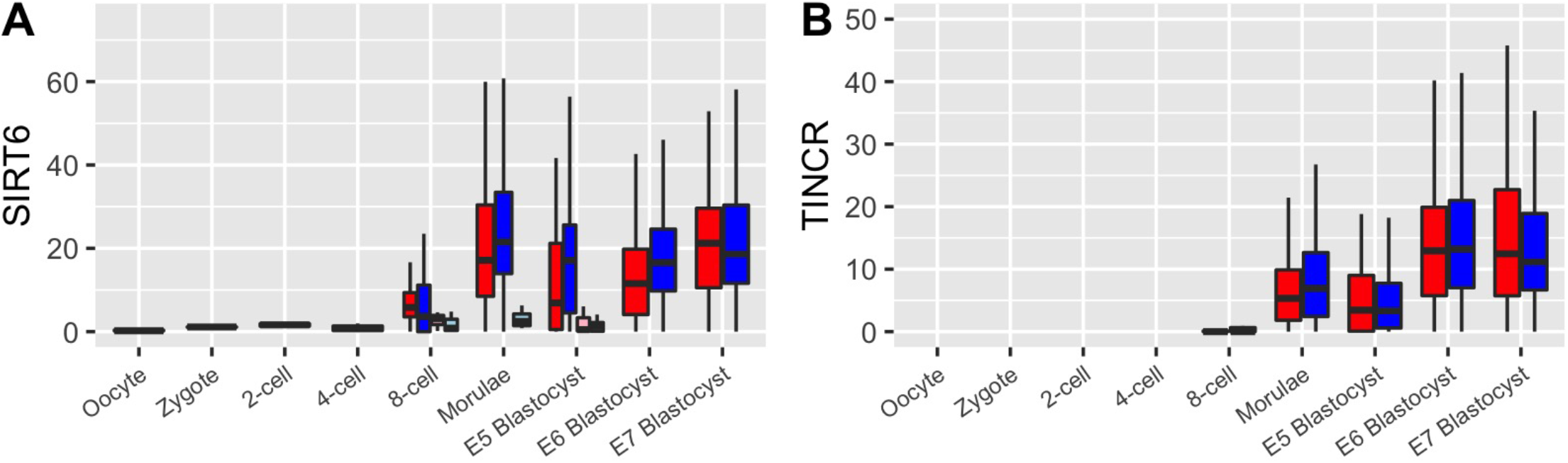
Box plots of gene expression (RPKM) determined by scRNA-seq in preimplantation human embryos for SIRT6 and TINCR.

**Supplemental Figure 2:**
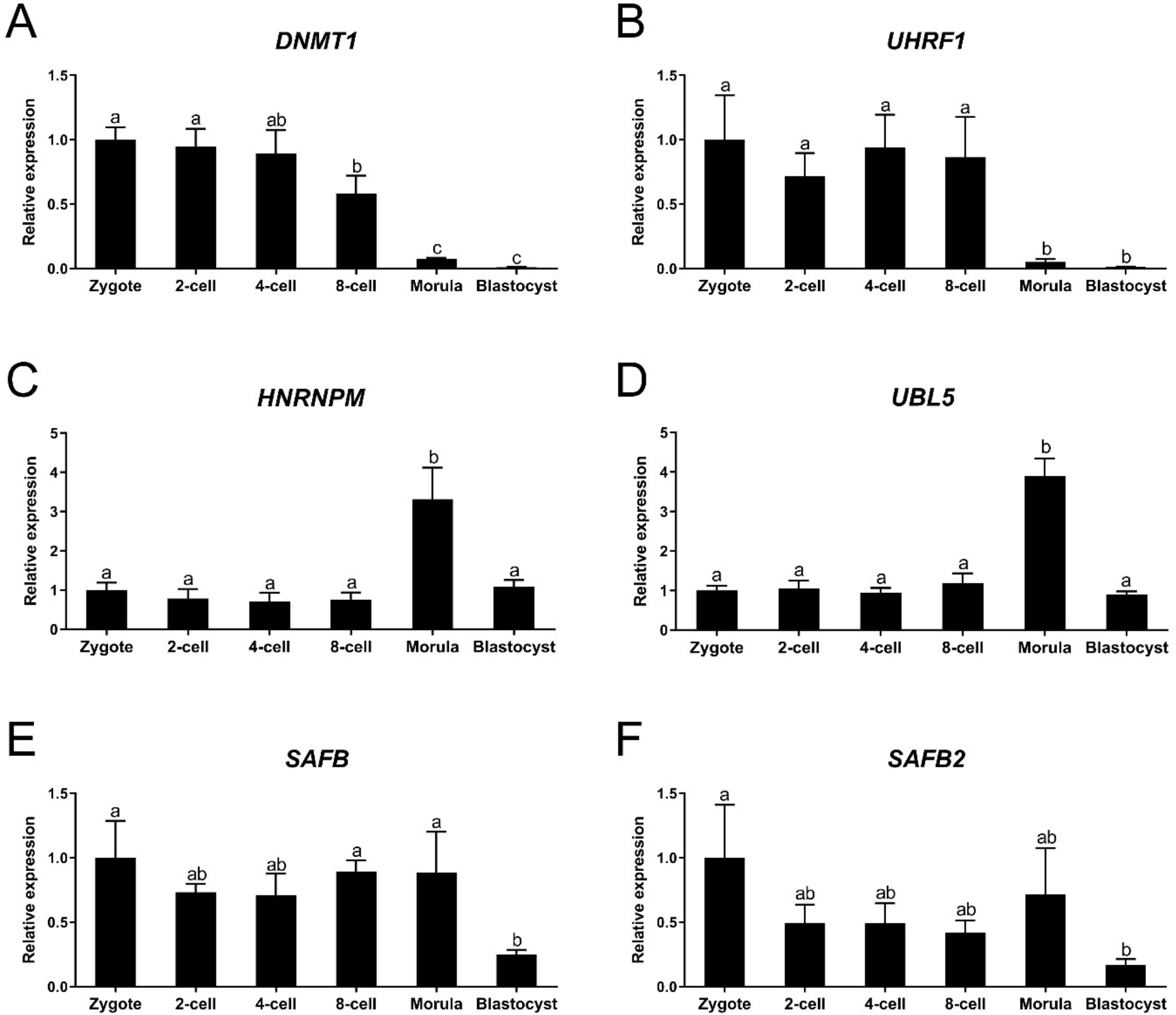
The relative expression of XCI related genes during in vitro bovine embryo development, as determined by quantitative RT-PCR. (A) DNMT1, (B) UHRF1, (C) HNRNPM, (D) UBL5, (E) SAFB, (F) SAFB2. Embryos were generated by fertilization with non-sexed sperm. Relative expression in GV oocytes is set at 1. Significant differences between bars are indicated by different letters above bars (p < 0.05). Error bars indicate standard deviations of three independent biological replicates.

